# Comparative analysis of flavivirus sfRNA dynamics and secondary structure

**DOI:** 10.64898/2026.04.07.716965

**Authors:** Camden R. Bair, David VanInsberghe, Anice C. Lowen, Graeme L. Conn, Christopher J. Neufeldt

## Abstract

The accumulation of subgenomic flavivirus RNAs (sfRNAs) modulates viral fitness and pathogenicity in culture and *in vivo*. These noncoding RNAs are produced by incomplete digestion of the flavivirus genome by the cellular 5’-3’ exoribonuclease (XRN1). Diverse flaviviruses have conserved RNA structural elements (RSEs) that map to their 3’-untranslated region (3’-UTR): Xrn-resistant RNA structures, dumbbell structures, and a 3’-stem loop (3’SL). Despite the importance of the 3’-UTR RSEs for flavivirus replication, the structural dynamics of sfRNA during flavivirus infection are understudied. Here, we use digital droplet PCR to quantify sfRNA levels during infection for a panel of mosquito-borne flaviviruses (MbFV) including dengue virus serotypes 1 (DENV1), 2 (DENV2), and 4 (DENV4), and Zika virus (ZIKV). We then used SHAPE-MaP on XRN1-digested, *in vitro*-transcribed sfRNAs from each virus to determine their secondary structures compared to the corresponding sfRNAs obtained from flavivirus-infected A549 cells. Results seen in-cell and *in vitro* were largely similar; however, motifs within the dumbbell, the small hairpin (sHP) directly upstream of the 3’-SL, and 3’-SL regions showed significant differences in the extent of nucleotide reactivity. These differences were consistent among the four flaviviruses examined and may indicate regions of sfRNA that are shielded by interaction with proteins or other nucleic acids during infection. However, strong protection indicative of sustained interaction was not apparent. Our findings suggest that sfRNA interactions with viral and host factors within the cell are few, occur via base-paired regions, or are highly transient.

**Importance:** Flaviviruses are highly prevalent human pathogens. The flavivirus genome contains RNA structural elements (RSEs), including those encoded in the 3’-UTR, that are necessary for viral replication. Subgenomic flavivirus RNAs (sfRNAs) are produced by incomplete digestion of flavivirus genomic RNA due to the cellular exoribonuclease XRN1 encountering 3’-UTR RSEs that promote its stalling and disassociation. Viruses unable to produce sfRNAs are highly attenuated, underlining their biological importance. sfRNA secondary structure has been investigated previously but little information is available on sfRNA secondary structure dynamics in infected cells. By comparing SHAPE-MaP reactivities *in vitro* and in cells, we determined that previously inferred structures are likely maintained within infected cells. We also identified differences in the extent of SHAPE reactivity between *in vitro* and in-cell environments that were common to multiple mosquito-borne flaviviruses. These differences suggest that sfRNAs may engage in transient interactions within the cell that may be important for their function.

## Introduction

Flaviviruses (genus *Orthoflavivirus*) are arthropod-borne viruses that pose a major global threat to human health (1). Infections by dengue virus alone cause an estimated 400 million symptomatic cases yearly, with endemic circulation in more than 100 countries across Latin

America, Africa, Asia, and Oceania (2). Other flaviviruses such as West Nile virus, Zika virus, Japanese encephalitis virus, and yellow fever virus also impact global health (3, 4). Owing to a lack of antivirals, limited availability or efficacy of vaccines, and shifts in vector ecology driven by climate change, flaviviruses continue to pose a serious threat to human health (5, 6).

Members of family *Flaviviridae* possess a ∼10kb positive-sense single-stranded RNA genome that contains a single open reading frame flanked by untranslated regions (UTRs) comprising conserved RNA structural elements (RSEs). During flavivirus replication, ∼0.5 kb subgenomic RNAs corresponding to the 3’-UTR are produced by incomplete digestion of the genome by the host 5’-3’ exoribonuclease enzyme XRN1 (7). XRN1 degrades genomic RNA from 5’ to 3’ but stalls and disassociates at one of the XRN-resistant structures (xrRNAs) located early in the 3’-UTR (7-9). This results in the accumulation of small non-coding RNAs comprising 3’-UTR RSEs (**Fig. 1A**). The accumulation of these small flavivirus RNAs (sfRNAs) enhances viral fitness in mammalian cells by inhibiting the interferon response (10, 11), preventing apoptosis (12), and counteracting RNAi (13). Furthermore, viruses unable to produce sfRNAs are highly attenuated in animal models (7, 14). While of long-standing interest, elucidating relationships between sfRNA structure and biological function in the native cellular context has only become more recently tractable with methodological advances in high throughput RNA probing strategies (15, 16).

**Figure 1:**
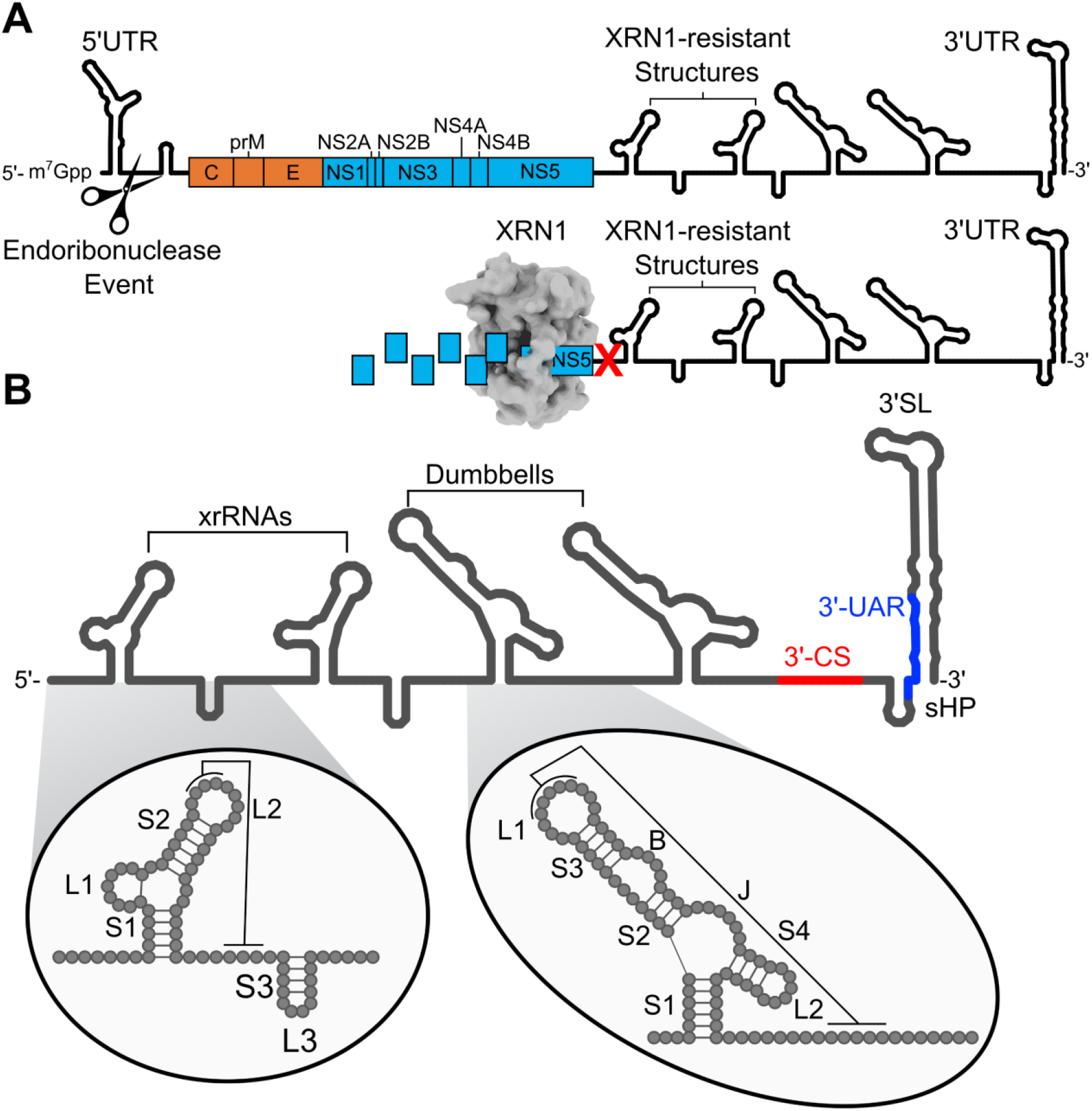
The highly structured flavivirus subgenomic RNAs (sfRNAs) are formed by incomplete digestion by XRN1. (A) Schematic of sfRNA biogenesis by the cellular XRN1 enzyme. Following an endonuclease event (depicted by scissors), XRN1 associates with monophosphorylated RNA and degrades gRNA from 5’ to 3’. XRN1 stalls and disassociates at xrRNA RSEs (red X). Cartoon illustrations depicting the 5’ and 3’ UTR flanking coding regions. Orange and blue ORFs indicate structural and nonstructural genes, respectively. Abbreviations: C,capsid; prM, pre-membrane; E, envelope; NS, nonstructural. B) Structure of flavivirus 3’-UTR/sfRNA RSEs. The 3’ cyclization (CS) and 3’ upstream AUG region (UAR) regions are highlighted in red and blue, respectively. Insets indicate labeling schemes for the xrRNA and DB structures. Abbreviations: S, stem; L, loop; B, bulge; J, junction.

Phylogenic analysis of flavivirus 3’-UTRs reveals multiple distinct clades with predicted 3’-UTR RNA structures within each clade characterized by conserved RSEs (17-21). For mosquito-borne flaviviruses (MbFVs), the 3’-UTR is composed of three primary RSEs: the xrRNA, dumbbell (DB), and terminal 3’-stem loop (3’SL) (**Fig. 1B**). Additionally, the 3’-UTR has been subject to duplication events leading to some viruses which contain multiple xrRNAs and DBs (19, 22). Duplication of these RSEs in some MbFVs appears to facilitate host adaptation and may have differing fitness effects depending on the host (19).

Selective 2′-hydroxyl acylation analyzed by primer extension (SHAPE) chemistries have been used to determine the secondary structures of full-length sfRNAs and isolated 3’-UTR RSEs for the prototypical flavivirus members (23-27). SHAPE reactivities have been reinforced by crystal structures of individual RSEs from various flaviviruses (20, 28-31). However, as these studies were largely performed outside of the context of infection, the 3’-UTR RNA secondary structure in the cellular environment remains understudied. Furthermore, given that sfRNAs and viral genomic RNA (gRNA) are localized in distinct cytoplasmic structures during infection (32), sfRNAs may possess differential structural dynamics than the gRNA 3’-UTR.

In this study, we examined the sfRNAs secondary structure dynamics for a panel of MbFVs. We applied SHAPE-MaP to XRN1-cleaved *in vitro*-transcribed RNA corresponding to the 3’-UTR and to sfRNAs from flavivirus-infected cells. We then used deltaSHAPE and Difference SHAPE calculations to compare the data obtained in each context. Our results do not support differences in RNA structure but rather reveal differences in the extent of SHAPE reactivity between *in vitro* and in-cell conditions that are consistent among MbFVs. These observations suggest possible sites for host or viral interactions that are transient in nature.

## Results

### sfRNA accumulation peaks 24 - 48 hours post infection

To define the sfRNA accumulation kinetics of MbFV, we applied digital droplet PCR to quantify gRNA and sfRNA during infection. We examined gRNA and sfRNA copy numbers from total RNA from infected A549 cells at 8, 18, 24, 36, and 48 hours post-infection (hpi) from a panel of MbFV, including the prototypical dengue virus serotype 2 (DENV2), dengue virus serotype 1 (DENV1), dengue virus serotype 4 (DENV4), and Zika virus (ZIKV). Genome copy number peaked at 18 hpi for DENV1, and ZIKV, while DENV2 and DENV4 lagged behind the other flaviviruses, with gRNA approaching peak accumulation at 24 hpi (**Fig. 2**). sfRNA accumulated more slowly than gRNA for DENV1, DENV4, and ZIKV, while sfRNA levels reached plateau at the same time point as gRNA for DENV2. sfRNA copy numbers were lower than gRNA levels for all viruses for the 8-, 18-, and 24-hour time points while sfRNA to gRNA copy numbers were comparable at later time points for DENV1, DENV4, and ZIKV. Notably, DENV2 sfRNA copy numbers never reached gRNA levels during the time course (**Fig. 2**). We observed peak sfRNA accumulation for all viruses between 24 and 48 hpi.

**Figure 2:**
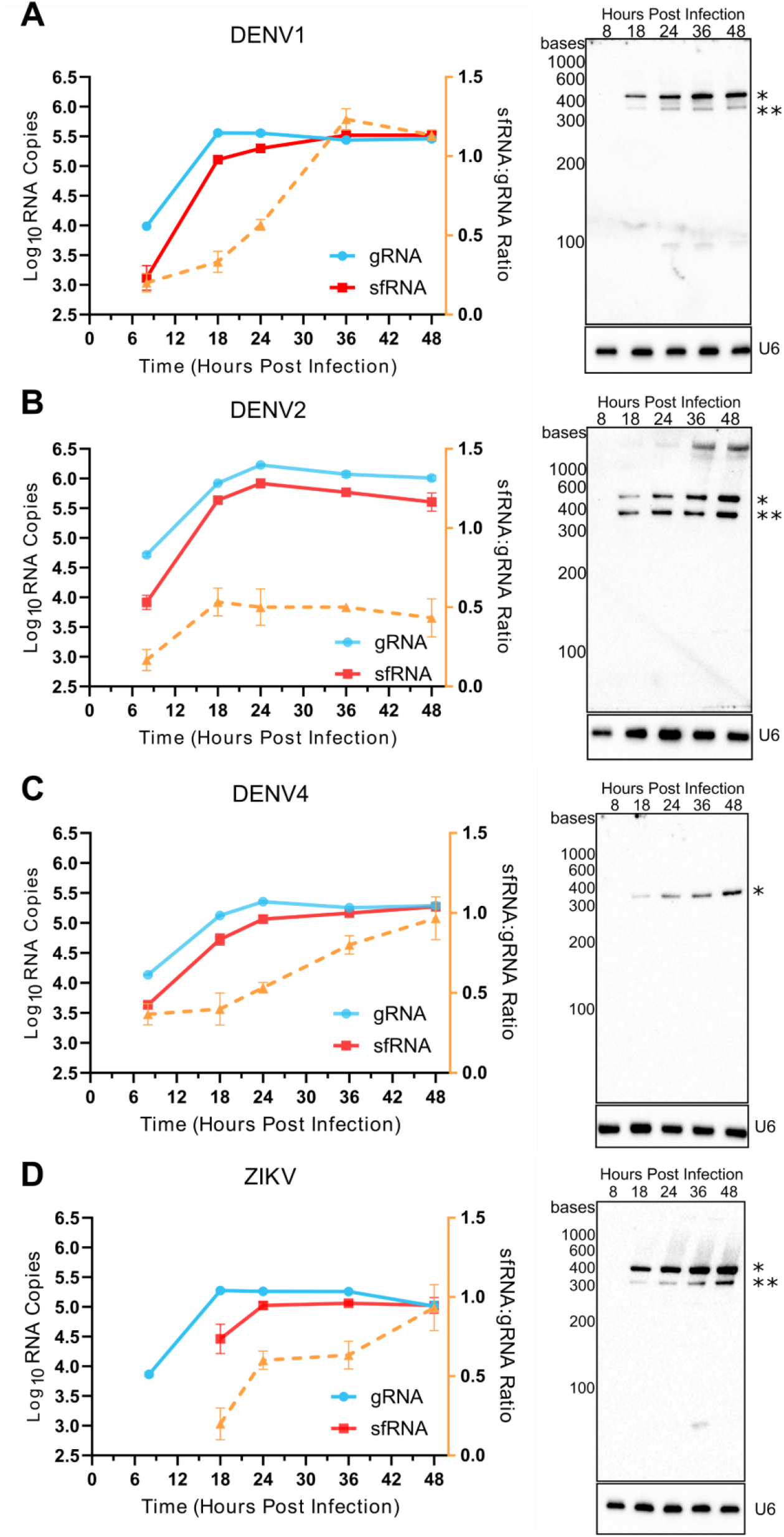
Flavivirus sfRNA accumulation peaks between 24 and 48 hpi. Digital droplet PCR quantification (left) of gRNA (blue), sfRNA (red), and the sfRNA:gRNA ratio (orange) for (A) DENV1-, (B) DENV2-, (C) DENV4- and (D) ZIKV-infected A549 cells. Data shown are mean ± SEM of three biological replicates. Northern blot analysis (right) of total RNA from flavivirus-infected A549 cells. sfRNA1(*) and sfRNA2(**) are indicated. The U6 snRNA was used as a loading control.

We also confirmed sfRNA accumulation using northern blot analysis and observed a similar trend of sfRNA reaching peak accumulation between 36 and 48 hpi. This approach also enabled visualization of distinct sfRNA species, including full-length sfRNAs (sfRNA1) and degradation intermediates (sfRNA2) which correspond to XRN1 stalling at the first or second xrRNA structure DENV1, DENV2, and ZIKV (**Fig. 2A-B, and D**). Only sfRNA1 was observed for DENV4 due to its single xrRNA (**Fig. 2C**). From these results we conclude that accumulation of gRNA and sfRNA are temporarily shifted during infection in A549 cells.

### sfRNA secondary structure in cells recapitulates secondary structure in vitro

To define the sfRNA secondary structural landscapes, we used SHAPE-MaP to compare *in vitro*-transcribed sfRNAs to sfRNAs from A549 cells infected with the same panel of MbFVs. These MbFVs represent variability in 3’-UTR RSE content: DENV1 and DENV2 possess two xrRNAs, two DBs, and the 3’-SL; DENV4 has the same complement of RSEs with the exception that it only carries one xrRNA; and, ZIKV, has two xrRNAs, one DB, a structure referred to as a pseudo-dumbbell, and the 3’-SL (18, 21). *In vitro* and in-cell SHAPE-MaP (33) analyses are shown for DENV4 in **Fig. 3** and the results for DENV1, DENV2, and ZIKV are reported in **Supplementary Fig. S2, S3, and S4**, respectively.

**Figure 3:**
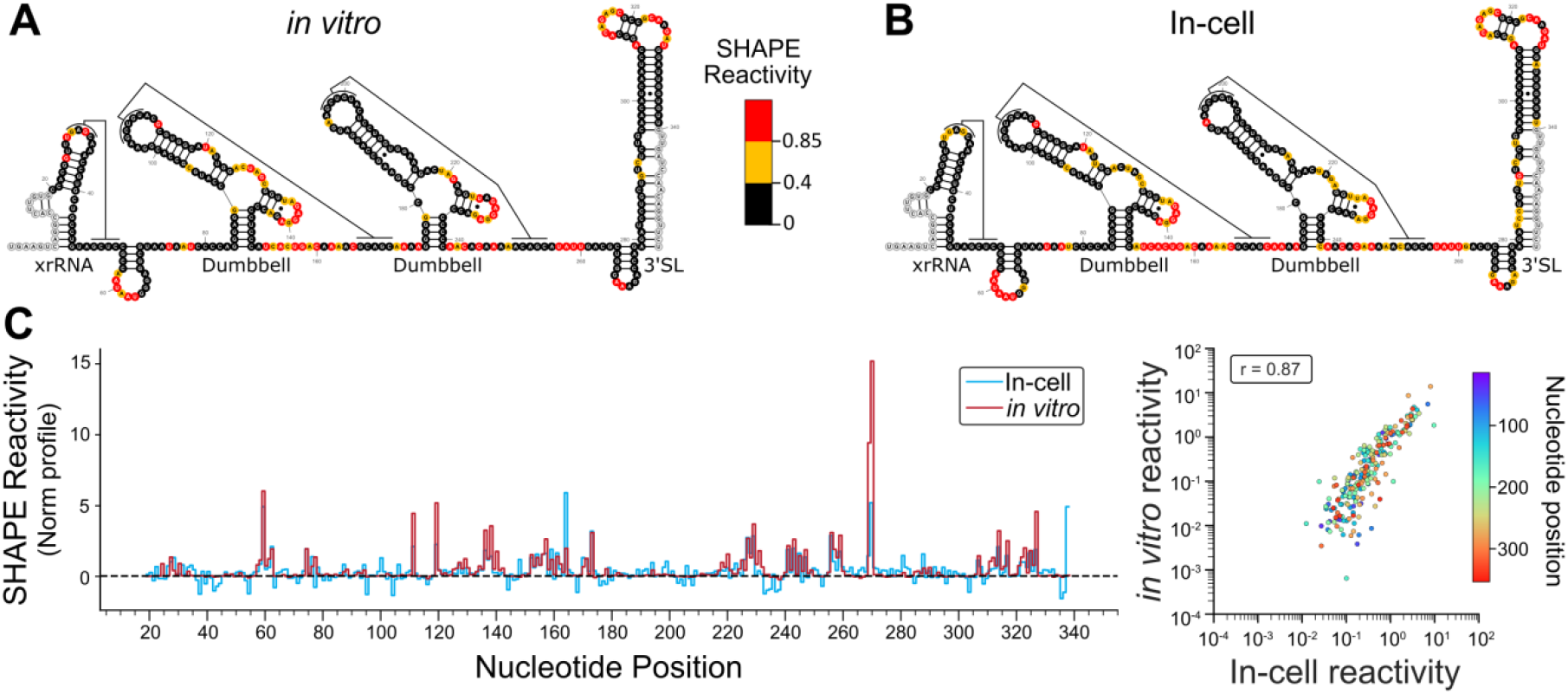
DENV4 sfRNA nucleotide SHAPE reactivity is highly similar *in vitro* and in virus-infected A549 cells. Normalized (A) *in vitro* and (B) in-cell SHAPE reactivity mapped to the predicted secondary structure of the DENV4 sfRNA. Predicted pseudoknot interactions in the xrRNA and DBs are indicated. Moderate and highly reactive nucleotides indicated by yellow and red fill, respectively. Undetermined SHAPE reactivity denoted by gray-outlined nucleotides. (C) Left: *in vitro* (red) and in-cell (blue) normalized SHAPE reactivity plotted against nucleotide position. Right: Pearson correlation analysis comparing *in vitro* and in-cell normalized SHAPE reactivity. Individual nucleotides are represented by a single point, and the color is indicative of the nucleotide position in the RNA. The Pearson correlation coefficient (r) is also reported.

We first performed SHAPE-MaP on *in vitro-*transcribed, XRN1-treated sfRNAs (**Supplementary Fig. S1**) and mapped normalized SHAPE reactivity to the predicted sfRNA secondary structures for DENV4, DENV1, DENV2, and ZIKV (**Fig. 3A** and **Supplementary Fig. S2A, S3A, and S4A**). The results for DENV2 and ZIKV compare well with previously published SHAPE reactivity data (23, 24). While equivalent analyses have not been performed previously for DENV1 and DENV4, the *in vitro* SHAPE reactivity of these sfRNAs also correlate well with that previously reported for DENV2. This outcome was expected based on sequence and structural conservation among MbFV RSEs. However, for DENV4, some novel features were apparent: we observed notable reactivity in the predicted xrRNA pseudoknot loop region (L2), indicating these interactions may be transient. Additionally, we observed low reactivity at the 3’ end of the DENV4 xrRNA L3 loop and the B region for the 3’-DB, indicating inaccessibility of these nucleotides to SHAPE reactivity due to additional nucleotide interactions or greater stability of these regions.

*In vitro* SHAPE probing does not consider the native cellular environment, in which RNA and/ or protein binding partners could alter the SHAPE reactivity observed for isolated sfRNAs. To examine potential differences due to the cellular environment, we therefore compared our *in vitro* results with SHAPE-probed sfRNAs extracted from flavivirus-infected A549 cells. Flavivirus-infected cells were treated with SHAPE reagent (2A3) at the time of peak sfRNA accumulation, as previously determined (**Fig. 2**). SHAPE-MaP was performed on sfRNAs and reactivity was mapped to the predicted secondary structure as before (**Fig. 3B** and **Supplementary Fig. S2B, S3B, and S4B**). Overall, SHAPE reactivity profiles of sfRNAs from infected cells correlated well with those for *in vitro* transcribed RNAs (**Fig. 3C** and **Supplementary Fig. S2C, S3C, and S4C**). Additionally, comparisons of mutation rates between *in vitro* and in-cell data sets were relatively consistent (**Supplementary Fig. S5**). Skyline plots comparing *in vitro* versus in-cell normalized reactivity reveal moderate per-nucleotide differences. However, despite these subtle differences for individual nucleotides, under both conditions, SHAPE reactivity was consistent with the predicted RSEs. Thus, in-cell SHAPE reactivities correlate well with those determined *in vitro* revealing that production of sfRNAs within cells does not impose durable changes on the formation of RSEs relative to those seen *in vitro*.

### Comparative analyses reveal changes in the extent of sfRNA SHAPE reactivity

To detect nuanced differences in mapped SHAPE reactivity between *in vitro* and in-cell conditions, we employed two established methods to analyze differences between these contexts: Difference SHAPE (34) and deltaSHAPE (35). Difference SHAPE reports increased or decreased in-cell reactivity at the single nucleotide level by subtracting the normalized *in vitro* SHAPE reactivity from the normalized in-cell reactivity. deltaSHAPE, meanwhile, analyzes statistical differences in SHAPE reactivity using a sliding window to identify regions of difference. As expected, these methodologies did not reveal regions of extensive structural difference between *in vitro* and in-cell derived sfRNAs. However, patterns of localized increased or decreased relative SHAPE reactivity were identified that were consistent across viruses.

Generally, in-cell reactivity was higher than that observed *in vitro* for the flexible nucleotide linkers between RSEs, particularly between the DBs for DENV1, DENV2, and DENV4 (**Fig. 4**). Additionally, Difference SHAPE revealed that the flexible nucleotides in the J region of the DBs had observable decreases in the extent of in-cell SHAPE reactivity. This was seen for all DBs except the DENV1 5’-DB. Consistent among all four viruses was a marked difference in the

**Figure 4:**
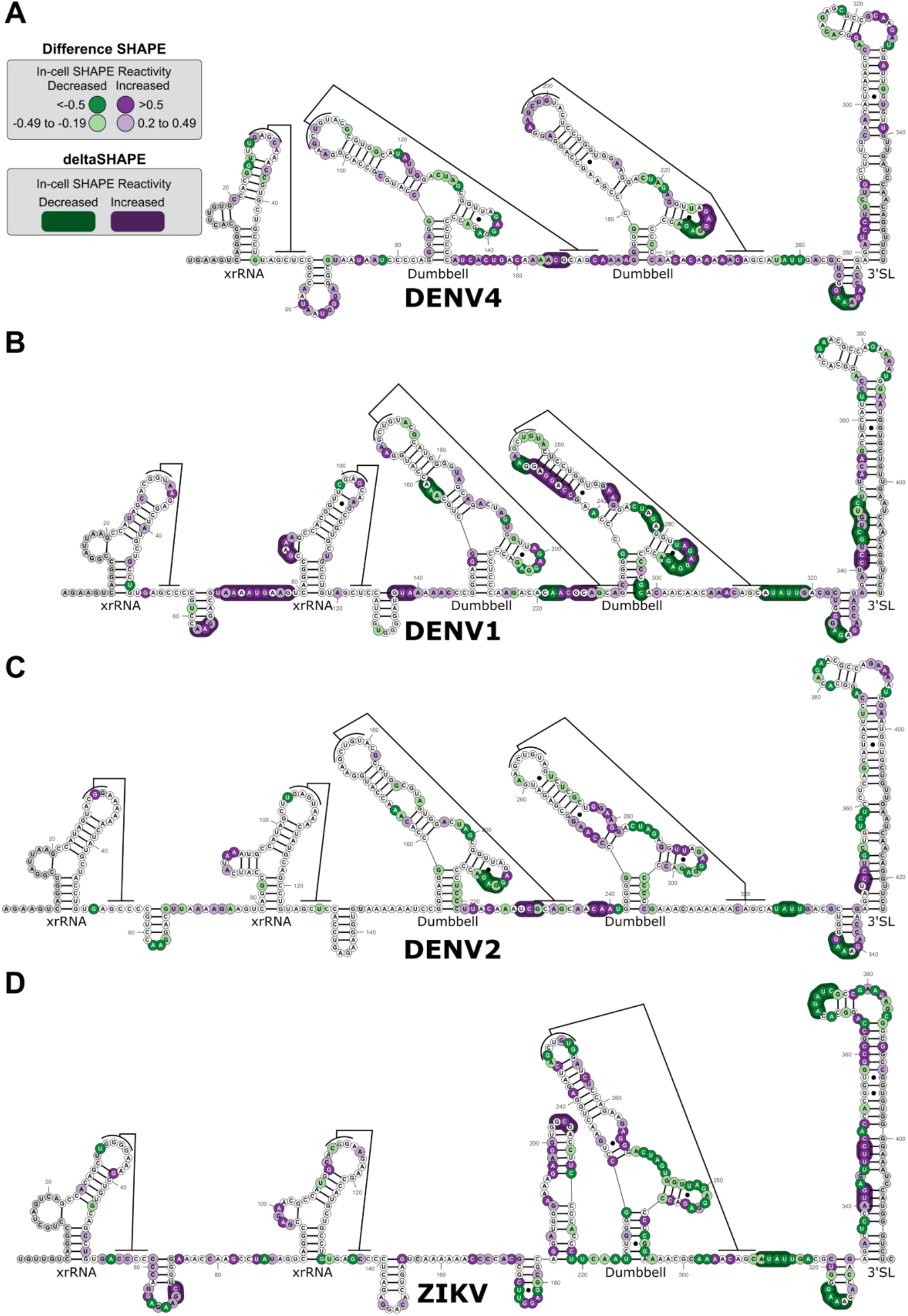
Difference SHAPE and deltaSHAPE analysis of flavivirus sfRNAs reveals consistent subtle differences between in vitro and in-cell SHAPE reactivity. Difference SHAPE and deltaSHAPE mapped to the predicted secondary structures of the (A) DENV4, (B) DENV1, (C) DENV2, and (D) ZIKV sfRNAs. Increased or decreased in-cell reactivity is annotated by filled circles in shades of purple (0.20– 0.49, or ≥0.50) or green (≤−0.50, or −0.49 to −0.19), respectively. No difference between *in vitro* and in-cell is denoted by white-filled nucleotides. Undetermined SHAPE reactivity are denoted by gray-filled nucleotides. In-cell enhancements and protections reported by deltaSHAPE aredenoted by outline shaded regions in dark purple and green, respectively.

SHAPE reactivity in the DB S4-L2 region. Specifically, decreased in-cell SHAPE reactivity was seen in this region and this difference was significant by deltaSHAPE for at least one DB for DENV1, DENV2, and DENV4. We also noted differences in SHAPE reactivity in the lower stem region of the 3’-SL. Finally, we observed significant protection in the small hairpin (sHP) and consistently decreased SHAPE reactivity in the unpaired nucleotides upstream of the sHP.

Together, these results reveal consistent, identifiable differences in the extent of in-cell SHAPE reactivity compared to *in vitro* across the MbFV panel with diverse RSE structural compositions. Importantly, these differences do not suggest large-scale changes in RSE structure. Rather, these differences indicate that the cellular environment gives rise to conditions that alter nucleotide modification by the SHAPE reagent, and may thus reflect sites of RNA-RNA, RNA-protein, or other interactions.

## Discussion

We investigated sfRNA production in infected cells and the sfRNA secondary structural dynamics using SHAPE-MaP under *in vitro* and in-cell conditions. We report sfRNA levels in A549 cells are generally less than gRNA levels, with sfRNA levels reaching plateau between 24 and 48 hpi. Our *in vitro* SHAPE-MaP results match those previously published for DENV2 and ZIKV and map well to the predicted secondary structure for DENV1 and DENV4. Similarly, SHAPE reactivity from flavivirus-infected A549 cells recapitulated the structural conservation with no major differences in RNA structure relative to *in vitro* conditions. Using two different SHAPE comparison methods, however, differences were observed in the extent of regions of local SHAPE reactivity, some of which were consistent among the viruses tested. These differences highlight the dynamic nature of these RNAs in cells, possibly due to inter-molecular interactions with RNAs or proteins. As such, this work provides a foundation for future exploration of the sfRNA function in the cellular environment.

Several reports have used northern blot analysis and quantitative PCR to quantify gRNA and sfRNA in infected cells or mosquitos (7, 10, 11, 14, 23, 29, 36-38). These reports demonstrate higher levels of sfRNAs compared to gRNAs with sfRNA:gRNA ratios ranging from 10:1 to >100:1. In contrast, we observed sfRNA:gRNA ratios ≤1.0. This difference may relate to our use of A549 cells, as prior reports probed RNA levels in infected mosquitos, mosquito cells, or mammalian cell lines such as BHK-21 or HUH-7 cells. Some viruses, such as Powassan virus, appear to produce considerably lower levels of sfRNA compared to gRNA (30). It is likely that the kinetics and efficiency of sfRNA production are dependent on the host system and/or viral species.

Flavivirus DBs are important structural elements of the flavivirus genome. Recent work has demonstrated roles for both Dengue virus DBs in viral replication within human and mosquito hosts, with the 3’-DB participating in long-range interactions with the 5’-UTR during genome cyclization (19). Flavivirus DBs show a remarkable level of structural conservation (31). Interestingly, the areas we found with consistent differences in SHAPE modification between *in vitro* and cellular environments (J and S4-L2; **Fig. 4**) are known to have high nucleotide conservation among MbFVs (31). Consistent with this evolutionary conservation, the relative nucleotide protection that we observed in cells may indicate that these regions are involved in weak or transient protein:RNA or RNA:RNA interactions during infection. Both viral and host proteins have been observed to bind viral 3’-UTR RSEs, but the specific binding sites for these proteins are unknown (39). Our comparative analyses showing consistent reactivity changes, particularly in the 3’-DB, may provide a framework for identifying protein binding sites within the 3’-UTR.

Genome cyclization during replication is accomplished predominantly via two long-range interactions between elements of the 5’-UTR and 3’-UTR: the upstream AUG region (UAR; a 5’-UTR sequence base-pairs with 3’-UTR nucleotides at the 3’ end of sHP and the 5’ portion of the 3’-SL) and the cyclization sequence (CS; the 5’-CS sequence interacts with the complimentary 3’-UTR sequence upstream of sHP) (40, 41). Disruption of these regions, including the sHP stem, alters the balance between linear and circular gRNA and reduces replication (42). Our structure probing demonstrated reactive nucleotides throughout the 3’-SL stem structure, particularly in the 3’-UAR (**Fig. 3**, and **Supplementary Fig. S2-S4**). This result recapitulated previous reports describing SHAPE probing of other flavivirus 3’-SL and our results are supported by the fact that this region, particularly the 3’-UAR, is conformationally flexible (43, 44). We also observed relative in-cell protection upstream of the sHP (3’-CS region), in the sHP loop, and in the 3’-UAR regions using both Difference SHAPE and deltaSHAPE (**Supplementary Fig. S4**). While the sHP loop is not predicted to form long-range interactions during cyclization, the observed enhanced and/or protected nucleotides in the CS and UAR regions do participate in complimentary base-paring with the 5’-UTR. These differences in the nucleotide SHAPE reactivity might be due to transient protein interactions taking place during infection or complimentary interactions with the 5’-UTR. One possible explanation might be physical interactions between sfRNAs and the 5’-UTR of genomic RNAs which would require further investigation.

There are some limitations to our study that are important to consider. First, although we enriched for sfRNAs to achieve at least a 10:1 ratio of sfRNA to genomes, full exclusion of gRNA from the in-cell preparations analyzed by SHAPE is not technically feasible. The gRNA 3’-UTR will therefore have made a modest contribution to the levels of SHAPE reactivity reported for in-cell samples. Second, we assume roughly equivalent nucleotide acetylation of the SHAPE reagent in both the *in vitro* and in-cell conditions. Limitations on the uptake of the SHAPE reagent by the cell would result in overall lower nucleotide acetylation in cells compared to *in vitro*. This phenomenon has been observed for some SHAPE reagents (45). To mitigate this concern, we chose the recently described 2A3 reagent because it is highly effective in various mammalian and prokaryotic cells (46).

Interventions that limit the expression or accumulation of sfRNAs during infection would be valuable. Live vaccine candidates that achieve attenuation through mutation to the 3’-UTR, specifically through deletion of portions of the 3’-DB, are currently in various stages of development (47-49). Construction of live flavivirus vaccines with blocked or attenuated sfRNA expression could provide promising candidates for future flavivirus vaccine development (50). Additionally, due to their conservation, the 3’-UTR RSEs are good candidates for therapeutics and antisense oligonucleotides targeting the 5’- and 3’-UTRs have shown promise to limit flavivirus infection in cells and in animals (51, 52). Flavivirus RSEs could also be targeted by small molecules to disrupt RNA structures (e.g., xrRNA, DB) to limit sfRNA production or reduce replication during infection. To capitalize on these opportunities, deeper understanding of the flavivirus 3’-UTR RSE structural and interaction dynamics and their roles during infection is needed. Our work confirms that sfRNAs fold as expected in cells during infection and provides a foundation for future studies investigating the function of sfRNAs.

## Materials and Methods

### Cells

A549 Cells (CCL-185) were obtained from the American Tissue Culture Collection. Cells were verified to be mycoplasma free using the PlasmoTest Detection kit (Invivogen, rep-pt1) and were maintained in DMEM (Gibco 11995073) supplemented with 10% fetal bovine serum (Gemeni R&D Systems Catalog number, S11550) and Normocin (Invivogen, ant-nr-2).

### Viruses

Molecular clones of DENV2 (strain 16681) and ZIKV (strain H/PF/2013) have been previously described and were a gracious gift of Ralf Bartenschlager (53, 54). Isolates of WHO reference strains for DENV1 (strain DV1-WP75) and DENV4 (strain DV4-H24) were generously provided by Dr. Matthew Collins (Emory University). For molecular clones, virus stocks were prepared as previously described by electroporation of Vero E6 cells with *in vitro* transcribed viral RNA; harvesting occurred 4-8 days after electroporation (55). For amplification of isolates or stocks from molecular clones, VeroE6 or C6/36 cells were infected at an multiplicity of infection (MOI) of 0.01 plaque-forming unit (PFU)/ml and supernatants were harvested 2-8 days after infection, beginning when cells showed cytopathic effect. Extracellular virus titers were determined by plaque assay in Vero E6 cells using a minimum essential medium (Gibco, 41090036) overlay containing 1.0% carboxymethylcellulose.

### Virus infections and RNA extraction

A549 cells were seeded in 12 well plates at a density of 3×10^5^ cells/well. Twenty-four hours post-seeding, cell culture medium was removed cells were infected with diluted virus (200 µl/well) at a MOI of 2 PFU/cell for 1 hour at 37°C with continuous rocking. Supernatant was removed and replaced with complete growth medium (1 ml/well). At 8, 12, 24, 36, and 48 hpi, growth media was removed and cells were lysed with buffer RLT from the RNeasy Plus RNA isolation kit (Qiagen, 74136) and frozen at -80°C. All lysates were thawed and RNA was extracted according to the manufacturer’s protocol.

### Northern blot analysis

Total RNA (1µg) was separated on a 6% denaturing polyacrylamide gel (Invitrogen, EC6865BOX). Gels were stained in RedSafe (iNtRON Biotechnology, 21141) to visualize RiboRuler Low Range ssRNA marker (Thermo Scientific, SM1831). RNA was transferred to Brightstar-Plus Positively Charged Nylon membrane (Invitrogen, AM10102) at 300 mA for 35 minutes in 0.5x TBE in an Owl Hep1 Electroblotter (Thermo Scientific, HEP-1). RNA was crosslinked in a UV Stratalinker 1800 (Agilent) using the auto setting. Membranes were prehybridized in ULTRAhub Oligo Hybridization buffer (Invitrogen, AM8663) at 42 °C for 30 minutes in a Shake ‘n’ Stack Hybridization Incubator (Thermo, Scientific, 6243). RNA was then probed with 5’-biotinylated DNA oligonucleotides (see **Supplementary Table S1** for probe sequences) at a final concentration of 5 nM in ULTRAhub Oligo Hybridization buffer overnight at 42 °C. Blots were washed in low stringency wash buffer (2x SSC (Invitrogen, 15557044), 0.1% SDS (MiliporeSigma, L3771)) three times at room temperature for 5 minutes each and then once at 42 °C for 30 minutes. Blots were then developed using the Chemiluminescent Nucleic Acid Detection Module (Thermo Scientific, 89880) following the manufacturer’s recommendations and imaged on a ChemiDoc MP imager (Bio-Rad). The imaged blot aligned with the RedSafe-stained gel image to obtain appropriate sizing of bands.

Blots were striped by incubating the blot in a minimal amount of boiling stripping solution (0.1X SSC, 0.5% SDS) for 15 minutes at room temperature. The blot was then hybridized, probed (with U6 loading control), and developed as described above.

### Reverse-transcription and sfRNA quantification by digital droplet PCR

Total RNA (1ug) was reverse-transcribed in 20 µl reactions using Maxima Reverse Transcriptase (Thermo Scientific, EP0743) with the provided buffer, Ribolock RNase inhibitor (Thermo Scientific, EO0381), dNTPs at a final concentration of 0.5 mM (Agilent, 200415-51), and virus-specific reverse-transcription primer (**Supplementary Table S1**). cDNAs were diluted 1:100 with water and used as a template for PCR using the QX200 ddPCR EvaGreen Supermix (Bio-Rad, 1864034) and primers anchored in the NS5 or 3’-UTR regions (**Supplementary Table S1**). Droplets were generated using a QX200 Droplet Generator (Bio-Rad, 1864002) and subsequently amplified in a thermocycler using the following conditions: 95 °C for 5 minutes; 40 cycles of 95 °C for 30 s, 58 °C for 1 minute; 4 °C for 5 minutes; 90 °C for 5 minutes; and hold at 4 °C. Positive droplets were then detected on the QX200 Droplet Reader (Bio-Rad, 1864003). gRNA copies were determined by the positive droplets from the NS5 primer set. sfRNA copies were calculated by subtracting the NS5 copies from the 3’-UTR copies.

### RNA synthesis

Plasmids encoding the sfRNAs from DENV1 (Western Pacific Strain (AY145121), nucleotides 10310-10736), DENV2 (Strain 16681 (NC_001474.2), nucleotides 10282-10723), DENV4 (TVP/360 strain (KU513442.1), nucleotides 10269-10649), and ZIKV (H/PF/2013 strain (KJ776791.2), nucleotides 10373-10807) were purchased from Genscript. sfRNA encoding-sequences were subcloned into an *in vitro* transcription plasmid encoding a 3’ Hepatitis Delta Virus ribozyme to ensure production of RNAs with homogenous 3’ ends (56). Plasmids were purified as previously described with the exception that ribosomal RNA was removed using Capto Core 700 multimodal chromatography resin (Cytiva, 17548102) (57). Plasmid was linearized using the EcoRV restriction enzyme (New England Biolabs, R3195T) and verified by agarose gel electrophoresis. Typical *in vitro* transcriptions were performed using 200 mM HEPES-KOH (pH 7.5), 28 mM MgCl_2_, 2 mM spermidine, 40 mM DTT, 10 mM rNTPs, 100 µg/ml of linearized template DNA, and 2.5 µl of T7 RNA polymerase (prepared in-house). Reactions were incubated at 37 °C for 3 hours (58). Pyrophosphate-magnesium precipitate was cleared using 0.2x volume of 250 mM EDTA (pH 8.0). RNA was dialyzed overnight at 4°C in 1x TE buffer (10 mM Tris-HCl (pH 8.0) and 1 mM EDTA) and ethanol precipitated. RNAs were separated using 6% native polyacrylamide gels (Invitrogen, EC6261BOX). RNA bands were visualized using UV shadowing at 254 nm, excised, and eluted using the crush-soak method in 0.3 M sodium acetate (Invitrogen, AM9740) followed by ethanol precipitation. RNA pellets were resuspended in 1x TE and the concentrations were determined by spectrophotometry.

### 2A3 synthesis

In-house synthesis of 2A3 was performed essentially as previously described (46). Briefly, 552.5 mg of 2-aminopyridine-3-carboxylic acid (MilliporeSigma, A68300) and 648.6 mg of 1,1′-carbonyldiimidazole (MilliporeSigma, 21860) were added to a 5 ml beaker. Anhydrous DMSO (4 ml, MilliporeSigma, 276855) was added and stirred at high speed continuously for 1 hour at room temperature. The solution was then spun at 16,100 x *g* for 2 minutes. The supernatant was transferred to cryogenic vials and stored at -80 °C until use.

### XRN1 treatment of in vitro-transcribed sfRNAs

Flavivirus RNA (2 µg) was heated to 95 °C for 2 minutes then snap cooled on ice. Concentrated folding buffer (final 50 mM Tris HCl pH 7.9, 100 mM NaCl, 10 mM MgCl_2_, 1 mM DTT) was then added and the RNA was heated to 55 °C for 10 minutes, slow cooled to 25 °C at 1 °C /min, and then chilled on ice. XRN1 (New England Biolabs, M0338, 2 units) and RppH (New England Biolabs, M0356,10 units) were then added and incubated at 37 °C for 1 hour. XRN1 cleavage was verified by electrophoresis on a 6% TBE-urea polyacrylamide gel (Invitrogen, EC6865BOX) stained with RedSafe.

### 2A3 treatment of in vitro-transcribed RNA

XRN1-digested RNA (20 µl total) was incubated at 37 °C for 5 minutes. RNA (10 µl) was then DMSO- or 2A3-treated (1 µl) and incubated at 37 °C for 15 minutes. The reaction was quenched by 10 µl of 1 M DTT. RNAs were then purified using Illustra MicroSpin G-25 Columns (Cytiva, 27-5325-01) following the manufacturer’s protocol.

### In-cell 2A3 treatment

A549 cells were flavivirus-infected at an MOI of 5 PFU/cell. 2A3 treatment was performed at 24 hpi (ZIKV) or 48 hpi (DENV1, DENV2, and DENV4). Cell culture medium was removed, and cells were washed once with 1x PBS. Prewarmed complete cell culture medium (900 µl) mixed with DMSO or 2A3 (100 µl) was used to treat cells for 15 minutes at 37 °C then neutralized with 1 ml of 1 M DTT. Total RNA was then extracted using TRIzol reagent (Invitrogen, 15596026).

### Enrichment of flavivirus sfRNA

Ribosomal RNA was removed from 5 µg of TRIzol-extracted total RNA using the RiboMinus Eukaryotic System v2 (Invitrogen, A15026) following the manufacturer’s protocol. RNA was then size separated using RNAClean XP beads (Beckman Coulter, A63987). Briefly, concentrated denaturing buffer (final 10 mM TrisHCl pH 8.0, 2 M urea) was combined with ribosomal RNA-depleted RNA to a total volume of 50 µl then heated to 90 °C for 5 minutes and snap cooled on ice. RNA was then separated with 25 µl (0.5x) RNAClean XP beads. The eluted RNA (Elution 1) was saved and the supernatant (RNA not bound to beads) was extracted again using 0.5x RNAClean XP beads. The eluted RNA (Elution 2) was saved, and the supernatant was extracted using 1.8x RNAClean XP beads. The eluted RNA (Elution 3) was saved. Each elution was evaluated for sfRNA enrichment using the digital droplet PCR assay previously discussed. Samples were processed for SHAPE-MaP if the sfRNA:gRNA enrichment was >10x.

### SHAPE-MaP library preparation

SHAPE-MaP was performed essentially as previously described (33). DMSO or 2A3-treated RNA was treated with 4 units of TURBO DNase (Invitrogen, AM2238) with the included buffer for 1 hour at 37 °C. DNase-treated RNA was then purified using the Monarch Spin RNA Cleanup kit (New England Biolabs, T2030L). RNA was eluted in 13 µl of nuclease-free water. RNA (10 µl) was then combined with 1 µl of 2 µM reverse-transcription primer (**Supplementary Table S1**), heated to 65 °C for 5 minutes, then immediately placed on ice. MaP buffer (33) and SuperScript II Reverse Transcriptase (Invitrogen, 18064014) were then added, mixed, and incubated at 37 °C for 3 hours. The reverse transcriptase was then heat inactivated at 70 °C for 5 minutes and the cDNA products were purified using Illustra MicroSpin G-25 Columns. Library preparation was performed by first PCR amplifying the sfRNA using Phusion High-fidelity DNA Polymerase (New England Biolabs, M0530L) and flavivirus-specific primers (**Supplementary Table S1**) and 5 µl of cDNA template in a 50 µl volume. PCR amplicons (5 µl) were electrophoresed on a 1% agarose TAE gel and products were visualized using RedSafe nucleic acid staining solution. Libraries were constructed with the Nextera XT DNA Library Preparation Kit (Illumina, FC-131-1096) and Illumina DNA/RNA UD Indexes with 1 ng cDNA (tagmented) as input. The final amplified libraries were purified using AMPure XP bead cleanup (Beckman Coulter, A63882). Libraries were validated by capillary electrophoresis on a TapeStation 4200 (Agilent), pooled at equimolar concentrations, and sequenced with PE100 reads on an NovaSeq 6000 (Illumina), targeting a depth of 500,000 reads per sample.

### Data analysis

Raw sequencing FASTQ files were inputted into the ShapeMapper2 (33, 59) for read alignment, mutation counting, and calculation of normalized nucleotide reactivity using default parameters with a minimum read depth of 5000. All ShapeMapper2 runs passed the built-in quality control checks. Linear regression of replicate data sets was performed using the RNAvigate pipeline (60). Comparison of per-nucleotide data was visualized using Skyline plots generated using RNAvigate. Normalized nucleotide reactivity was mapped to the predicted sfRNA secondary structures using RNAcanvas (61). Comparison of *in vitro* and in-cell SHAPE-MaP reactivity was performed using Difference SHAPE (34) and deltaSHAPE (35). The per-nucleotide averaged *in vitro* normalized SHAPE reactivities were subtracted from the in-cell data set. The difference in in-cell reactivity for each nucleotide was then classified as decreased (≤-0.50, or -0.49 to -0.19), no change (-0.20 to 0.19), or increased (0.20–0.49, or ≥0.50). For deltaSHAPE, SHAPE-MaP-generated profiles.txt files were averaged between replicates then used as raw data for the execution of the deltaSHAPE pipeline integrated into the RNAvigate suite using default parameters. Enhanced and protected nucleotides were then mapped to the secondary structure of each sfRNA using RNAvigate and transferred to RNAcanvas. ShapeMapper2 mutation rates were plotted using GraphPad Prism version 11.0.0 for Windows (www.graphpad.com).

## Data Availability

Sequences are available through the NCBI Sequence Read Archive (https://www.ncbi.nlm.nih.gov/sra) BioProject accession number PRJNA1443375.

## Acknowledgements

This work was supported by the National Institute of Allergy and Infectious Disease through awards R01AI144067 to GLC and R01AI185849 to CJN. Next generation sequencing services were provided by the Emory NPRC Genomics Core (RRID:SCR_026418) which is supported in part by NIH P51OD011132. Sequencing data was acquired on an Illumina NovaSeq 6000 funded by NIH S10OD026799. We also thank Dr. Danny Incarnato for helpful discussion on 2A3 synthesis.

## References

1. Pierson TC, Diamond MS. 2020. The continued threat of emerging flaviviruses. Nat Microbiol 5:796–812.

2. Bhatt S, Gething PW, Brady OJ, Messina JP, Farlow AW, Moyes CL, Drake JM, Brownstein JS, Hoen AG, Sankoh O, Myers MF, George DB, Jaenisch T, Wint GR, Simmons CP, Scott TW, Farrar JJ, Hay SI. 2013. The global distribution and burden of dengue. Nature 496:504–7.

3. Pierson TC, Diamond MS. 2018. The emergence of Zika virus and its new clinical syndromes. Nature 560:573–581.

4. Roehrig JT. 2013. West nile virus in the United States - a historical perspective. Viruses 5:3088–108.

5. Dutta SK, Langenburg T. 2023. A Perspective on Current Flavivirus Vaccine Development: A Brief Review. Viruses 15.

6. Bifani AM, Chan KWK, Borrenberghs D, Tan MJA, Phoo WW, Watanabe S, Goethals O, Vasudevan SG, Choy MM. 2023. Therapeutics for flaviviral infections. Antiviral Res 210:105517.

7. Pijlman GP, Funk A, Kondratieva N, Leung J, Torres S, van der Aa L, Liu WJ, Palmenberg AC, Shi PY, Hall RA, Khromykh AA. 2008. A highly structured, nuclease-resistant, noncoding RNA produced by flaviviruses is required for pathogenicity. Cell Host Microbe 4:579–91.

8. Chang JH, Xiang S, Xiang K, Manley JL, Tong L. 2011. Structural and biochemical studies of the 5’-->3’ exoribonuclease Xrn1. Nat Struct Mol Biol 18:270–6.

9. Clarke BD, Roby JA, Slonchak A, Khromykh AA. 2015. Functional non-coding RNAs derived from the flavivirus 3’ untranslated region. Virus Res 206:53–61.

10. Manokaran G, Finol E, Wang C, Gunaratne J, Bahl J, Ong EZ, Tan HC, Sessions OM, Ward AM, Gubler DJ, Harris E, Garcia-Blanco MA, Ooi EE. 2015. Dengue subgenomic RNA binds TRIM25 to inhibit interferon expression for epidemiological fitness. Science 350:217–21.

11. Bidet K, Dadlani D, Garcia-Blanco MA. 2014. G3BP1, G3BP2 and CAPRIN1 are required for translation of interferon stimulated mRNAs and are targeted by a dengue virus non-coding RNA. PLoS Pathog 10:e1004242.

12. Slonchak A, Hugo LE, Freney ME, Hall-Mendelin S, Amarilla AA, Torres FJ, Setoh YX, Peng NYG, Sng JDJ, Hall RA, van den Hurk AF, Devine GJ, Khromykh AA. 2020. Zika virus noncoding RNA suppresses apoptosis and is required for virus transmission by mosquitoes. Nat Commun 11:2205.

13. Schnettler E, Sterken MG, Leung JY, Metz SW, Geertsema C, Goldbach RW, Vlak JM, Kohl A, Khromykh AA, Pijlman GP. 2012. Noncoding flavivirus RNA displays RNA interference suppressor activity in insect and Mammalian cells. J Virol 86:13486–500.

14. Funk A, Truong K, Nagasaki T, Torres S, Floden N, Balmori Melian E, Edmonds J, Dong H, Shi PY, Khromykh AA. 2010. RNA structures required for production of subgenomic flavivirus RNA. J Virol 84:11407–17.

15. Cao X, Zhang Y, Ding Y, Wan Y. 2024. Identification of RNA structures and their roles in RNA functions. Nat Rev Mol Cell Biol 25:784–801.

16. Wang XW, Liu CX, Chen LL, Zhang QC. 2021. RNA structure probing uncovers RNA structure-dependent biological functions. Nat Chem Biol 17:755–766.

17. Ochsenreiter R, Hofacker IL, Wolfinger MT. 2019. Functional RNA Structures in the 3’UTR of Tick-Borne, Insect-Specific and No-Known-Vector Flaviviruses. Viruses 11.

18. Villordo SM, Carballeda JM, Filomatori CV, Gamarnik AV. 2016. RNA Structure Duplications and Flavivirus Host Adaptation. Trends Microbiol 24:270–283.

19. de Borba L, Villordo SM, Marsico FL, Carballeda JM, Filomatori CV, Gebhard LG, Pallares HM, Lequime S, Lambrechts L, Sanchez Vargas I, Blair CD, Gamarnik AV. 2019. RNA Structure Duplication in the Dengue Virus 3’ UTR: Redundancy or Host Specificity? mBio 10.

20. Slonchak A, Parry R, Pullinger B, Sng JDJ, Wang X, Buck TF, Torres FJ, Harrison JJ, Colmant AMG, Hobson-Peters J, Hall RA, Tuplin A, Khromykh AA. 2022. Structural analysis of 3’UTRs in insect flaviviruses reveals novel determinants of sfRNA biogenesis and provides new insights into flavivirus evolution. Nat Commun 13:1279.

21. Finol E, Ooi EE. 2019. Evolution of Subgenomic RNA Shapes Dengue Virus Adaptation and Epidemiological Fitness. iScience 16:94–105.

22. Gritsun TS, Gould EA. 2006. Direct repeats in the 3’ untranslated regions of mosquito-borne flaviviruses: possible implications for virus transmission. J Gen Virol 87:3297– 3305.

23. Chapman EG, Moon SL, Wilusz J, Kieft JS. 2014. RNA structures that resist degradation by Xrn1 produce a pathogenic Dengue virus RNA. Elife 3:e01892.

24. Zhang Y, Zhang Y, Liu ZY, Cheng ML, Ma J, Wang Y, Qin CF, Fang X. 2019. Long non-coding subgenomic flavivirus RNAs have extended 3D structures and are flexible in solution. EMBO Rep 20:e47016.

25. Madden EA, Plante KS, Morrison CR, Kutchko KM, Sanders W, Long KM, Taft-Benz S, Cruz Cisneros MC, White AM, Sarkar S, Reynolds G, Vincent HA, Laederach A, Moorman NJ, Heise MT. 2020. Using SHAPE-MaP To Model RNA Secondary Structure and Identify 3’UTR Variation in Chikungunya Virus. J Virol 94.

26. Boerneke MA, Gokhale NS, Horner SM, Weeks KM. 2023. Structure-first identification of RNA elements that regulate dengue virus genome architecture and replication. Proc Natl Acad Sci U S A 120:e2217053120.

27. Li P, Wei Y, Mei M, Tang L, Sun L, Huang W, Zhou J, Zou C, Zhang S, Qin CF, Jiang T, Dai J, Tan X, Zhang QC. 2018. Integrative Analysis of Zika Virus Genome RNA Structure Reveals Critical Determinants of Viral Infectivity. Cell Host Microbe 24:875–886 e5.

28. Akiyama BM, Laurence HM, Massey AR, Costantino DA, Xie X, Yang Y, Shi PY, Nix JC, Beckham JD, Kieft JS. 2016. Zika virus produces noncoding RNAs using a multi-pseudoknot structure that confounds a cellular exonuclease. Science 354:1148–1152.

29. Chapman EG, Costantino DA, Rabe JL, Moon SL, Wilusz J, Nix JC, Kieft JS. 2014. The structural basis of pathogenic subgenomic flavivirus RNA (sfRNA) production. Science 344:307–10.

30. Langeberg CJ, Szucs MJ, Sherlock ME, Vicens Q, Kieft JS. 2025. Tick-borne flavivirus exoribonuclease-resistant RNAs contain a double loop structure. Nat Commun 16:4515.

31. Akiyama BM, Graham ME, Z OD, Beckham JD, Kieft JS. 2021. Three-dimensional structure of a flavivirus dumbbell RNA reveals molecular details of an RNA regulator of replication. Nucleic Acids Res 49:7122–7138.

32. Watkins JM, Burke JM. 2024. RNase L-induced bodies sequester subgenomic flavivirus RNAs to promote viral RNA decay. Cell Rep 43:114694.

33. Smola MJ, Rice GM, Busan S, Siegfried NA, Weeks KM. 2015. Selective 2’-hydroxyl acylation analyzed by primer extension and mutational profiling (SHAPE-MaP) for direct, versatile and accurate RNA structure analysis. Nat Protoc 10:1643–69.

34. Strassler SE, Bowles IE, Krishnamohan A, Kim H, Edgington CB, Kuiper EG, Hancock CJ, Comstock LR, Jackman JE, Conn GL. 2023. tRNA m(1)G9 modification depends on substrate-specific RNA conformational changes induced by the methyltransferase Trm10. J Biol Chem 299:105443.

35. Smola MJ, Calabrese JM, Weeks KM. 2015. Detection of RNA-Protein Interactions in Living Cells with SHAPE. Biochemistry 54:6867–75.

36. Syenina A, Vijaykrishna D, Gan ES, Tan HC, Choy MM, Siriphanitchakorn T, Cheng C, Vasudevan SG, Ooi EE. 2020. Positive epistasis between viral polymerase and the 3’ untranslated region of its genome reveals the epidemiologic fitness of dengue virus. Proc Natl Acad Sci U S A 117:11038–11047.

37. Pompon J, Manuel M, Ng GK, Wong B, Shan C, Manokaran G, Soto-Acosta R, Bradrick SS, Ooi EE, Misse D, Shi PY, Garcia-Blanco MA. 2017. Dengue subgenomic flaviviral RNA disrupts immunity in mosquito salivary glands to increase virus transmission. PLoS Pathog 13:e1006535.

38. Yeh SC, Diosa-Toro M, Tan WL, Rachenne F, Hain A, Yeo CPX, Bribes I, Xiang BWW, Sathiamoorthy Kannan G, Manuel MC, Misse D, Mok YK, Pompon J. 2022. Characterization of dengue virus 3’UTR RNA binding proteins in mosquitoes reveals that AeStaufen reduces subgenomic flaviviral RNA in saliva. PLoS Pathog 18:e1010427.

39. Roby JA, Pijlman GP, Wilusz J, Khromykh AA. 2014. Noncoding subgenomic flavivirus RNA: multiple functions in West Nile virus pathogenesis and modulation of host responses. Viruses 6:404–27.

40. Villordo SM, Gamarnik AV. 2009. Genome cyclization as strategy for flavivirus RNA replication. Virus Res 139:230–9.

41. Khromykh AA, Meka H, Guyatt KJ, Westaway EG. 2001. Essential role of cyclization sequences in flavivirus RNA replication. J Virol 75:6719–28.

42. Villordo SM, Alvarez DE, Gamarnik AV. 2010. A balance between circular and linear forms of the dengue virus genome is crucial for viral replication. RNA 16:2325–35.

43. Sztuba-Solinska J, Teramoto T, Rausch JW, Shapiro BA, Padmanabhan R, Le Grice SF. 2013. Structural complexity of Dengue virus untranslated regions: cis-acting RNA motifs and pseudoknot interactions modulating functionality of the viral genome. Nucleic Acids Res 41:5075–89.

44. Brinton MA, Basu M. 2015. Functions of the 3’ and 5’ genome RNA regions of members of the genus Flavivirus. Virus Res 206:108–19.

45. Busan S, Weidmann CA, Sengupta A, Weeks KM. 2019. Guidelines for SHAPE Reagent Choice and Detection Strategy for RNA Structure Probing Studies. Biochemistry 58:2655–2664.

46. Marinus T, Fessler AB, Ogle CA, Incarnato D. 2021. A novel SHAPE reagent enables the analysis of RNA structure in living cells with unprecedented accuracy. Nucleic Acids Res 49:e34.

47. Kallás EG, Cintra MAT, Moreira JA, Patiño EG, Braga PE, Tenório JCV, Infante V, Palacios R, de Lacerda MVG, Pereira DB, da Fonseca AJ, Gurgel RQ, Coelho ICB, Fontes CJF, Marques ETA, Romero GAS, Teixeira MM, Siqueira AM, Barral AMP, Boaventura VS, Ramos F, Elias E Jr, de Moraes JC, Covas DT, Kalil J, Precioso AR, Whitehead SS, Esteves-Jaramillo A, Shekar T, Lee JJ, Macey J, Kelner SG, Coller BAG, Boulos FC, Nogueira ML. 2024. Live, Attenuated, Tetravalent Butantan-Dengue Vaccine in Children and Adults. New England Journal of Medicine 390:397–408.

48. Russell KL, Rupp RE, Morales-Ramirez JO, Diaz-Perez C, Andrews CP, Lee AW, Finn TS, Cox KS, Falk Russell A, Schaller MM, Martin JC, Hyatt DM, Gozlan-Kelner S, Bili A, Coller BG. 2022. A phase I randomized, double-blind, placebo-controlled study to evaluate the safety, tolerability, and immunogenicity of a live-attenuated quadrivalent dengue vaccine in flavivirus-naive and flavivirus-experienced healthy adults. Hum Vaccin Immunother 18:2046960.

49. Whitehead SS, Durbin AP, Pierce KK, Elwood D, McElvany BD, Fraser EA, Carmolli MP, Tibery CM, Hynes NA, Jo M, Lovchik JM, Larsson CJ, Doty EA, Dickson DM, Luke CJ, Subbarao K, Diehl SA, Kirkpatrick BD. 2017. In a randomized trial, the live attenuated tetravalent dengue vaccine TV003 is well-tolerated and highly immunogenic in subjects with flavivirus exposure prior to vaccination. PLoS Negl Trop Dis 11:e0005584.

50. Doets K, Pijlman GP. 2024. Subgenomic flavivirus RNA as key target for live-attenuated vaccine development. J Virol 98:e0010023.

51. Deas TS, Bennett CJ, Jones SA, Tilgner M, Ren P, Behr MJ, Stein DA, Iversen PL, Kramer LD, Bernard KA, Shi PY. 2007. In vitro resistance selection and in vivo efficacy of morpholino oligomers against West Nile virus. Antimicrob Agents Chemother 51:2470– 82.

52. Kinney RM, Huang CY, Rose BC, Kroeker AD, Dreher TW, Iversen PL, Stein DA. 2005. Inhibition of dengue virus serotypes 1 to 4 in vero cell cultures with morpholino oligomers. J Virol 79:5116–28.

53. Munster M, Plaszczyca A, Cortese M, Neufeldt CJ, Goellner S, Long G, Bartenschlager R. 2018. A Reverse Genetics System for Zika Virus Based on a Simple Molecular Cloning Strategy. Viruses 10.

54. Fischl W, Bartenschlager R. 2013. High-throughput screening using dengue virus reporter genomes. Methods Mol Biol 1030:205–19.

55. Owen JE, Bemis CL, Wang Q, Varadan AC, Vander Velden JW, Andacic L, Uyar O, Gupta M, Scharer CD, Chatel-Chaix L, Scaturro P, Suthar MS, Neufeldt CJ. 2025. Homotypic endoplasmic reticulum membrane tethering is critical for flavivirus replication. bioRxiv doi:10.1101/2025.11.26.690702.

56. Walker SC, Avis JM, Conn GL. 2003. General plasmids for producing RNA in vitro transcripts with homogeneous ends. Nucleic Acids Res 31:e82.

57. Linpinsel JL, Conn GL. 2012. General protocols for preparation of plasmid DNA template, RNA in vitro transcription, and RNA purification by denaturing PAGE. Methods Mol Biol 941:43–58.

58. Gurevich VV. 1996. Use of bacteriophage RNA polymerase in RNA synthesis. Methods Enzymol 275:382–97.

59. Busan S, Weeks KM. 2018. Accurate detection of chemical modifications in RNA by mutational profiling (MaP) with ShapeMapper 2. RNA 24:143–148.

60. Irving PS, Weeks KM. 2024. RNAvigate: efficient exploration of RNA chemical probing datasets. Nucleic Acids Res 52:2231–2241.

61. Johnson PZ, Simon AE. 2023. RNAcanvas: interactive drawing and exploration of nucleic acid structures. Nucleic Acids Res 51:W501–W508.

